# CD81-guided heterologous EVs present heterogeneous interactions with breast cancer cells

**DOI:** 10.1101/2024.02.06.579138

**Authors:** Elena Gurrieri, Giulia Carradori, Michela Roccuzzo, Michael Pancher, Daniele Peroni, Romina Belli, Caterina Trevisan, Michela Notarangelo, Wen-Qiu Huang, Agata SA Carreira, Alessandro Quattrone, Guido Jenster, Timo L.M. Ten Hagen, Vito Giuseppe D’Agostino

## Abstract

Extracellular vesicles (EVs) are cell-secreted particles conceived as natural vehicles for intercellular communication. The intrinsic biocompatibility, stability in biofluids, and heterogeneous molecular cargo of EVs promise advancements in targeted therapy applications. However, predicting cell-targeting spectrum and cargo delivery are fundamental challenges for exploiting EVs or hybrid formulations. In this work, we combined cell-based and biochemical approaches to understand if secreted EVs show predictable EV-cell interactions and consequent cargo delivery. We exploited the tetraspanin CD81 to encode full-length recombinant proteins with a C-terminal GFP reporter encompassing or not Trastuzumab light chains targeting the HER2 receptor. These fusion proteins participated in vesicular trafficking dynamics and accumulated on secreted EVs when transiently over-expressed in HEK293T cells. Despite the presence of GFP, secreted EV populations retained a HER2 receptor-binding capacity and were used in EV-cell interaction assays. In time-frames where the global GFP spot distribution did not change between HER2-positive (SK-BR-3) or –negative (MDA-MB-231) breast cancer cell lines, the HER2 manipulation in isogenic cells remarkably affected the tropism of heterologous EVs. In this line, secreted doxorubicin-EVs, which showed improved efficacy compared to the free drug, had a reduced cell-killing activity on SK-BR-3 with a knocked-out HER2 receptor. Interestingly, the fusion protein-corresponding transcripts also present as full-length mRNAs in recombinant EVs could reach orthotopic breast tumors in JIMT-1-xenografted mice, as detected by ddPCR in tissue biopsies, improving our sensitivity in detecting bioavailable cargoes. These data show multiple mechanisms underlying EV-cell interactions and prioritize the profiling of surfaceomes for better comprehension of cell engagement and design new generations of EV-based nanovehicles.

## Introduction

Extracellular vesicles (EVs) are cell-secreted lipid particles recognized as natural delivery vehicles for intercellular communication(Théry et al., 2018), (Liu and Wang, 2023). The heterogeneous molecular cargo and the intrinsic biocompatibility and stability in biofluids (Herrmann et al., 2020) render EVs an attractive source for developing novel bioformulations for targeted therapy(Sarkar et al., 2023). EVs comprise vesicular particles deriving from different biogenesis pathways, i.e., exosomes and microvesicles, besides apoptotic bodies and non-vesicular structures called exomeres (Anand et al., 2021). Exosomes are small vesicles, typically 30 to 150 nm in diameter, formed within multivesicular bodies (MVBs). These particles are released in the extracellular space upon MVB fusion with the plasma membrane(Van Niel et al., 2018). Both endosomal sorting complex required for transport (ESCRT)-dependent or -independent mechanisms are involved in the formation of MVBs and intraluminal vesicles, including proteins such as ALIX (ALG2-interacting protein X), TSG101 (tumor susceptibility gene 101) or SYNTENIN (Baietti et al., 2012). Tetraspanins like CD81, CD63, and CD9 are described in the ESCRT-independent mechanism and form clusters with transmembrane/cytosolic proteins involved in membrane inward budding (Gurung et al., 2021). On the other hand, microvesicles or ectosomes are typically larger EVs ranging from 150 nm to 1 μm in diameter. These are generated by direct outward budding of the plasma membrane in response to signaling and membrane rearrangements, which are yet not fully clarified, including changes in lipid-protein composition and calcium-dependent mechanisms shaping the membrane symmetry in connection with the cytoskeleton (Van Niel et al., 2018).

Recent seminal studies highlighted the EV potential in delivering siRNAs, small molecules, and microRNAs (Möller and Lobb, 2020). Through surface engineering and cargo loading techniques, EVs were reported to exert anti-inflammatory and antitumor effects, or neuroprotection and tissue regeneration (Muhammad et al., 2023), (Roudi et al., 2023), (Greening et al., 2023).

The Human Epidermal Growth Factor Receptor 2 (HER2/ERBB2) is a tyrosine kinase member of the EGF receptor family (Abunada et al., 2023). Aberrant protein expression and mutations in the *ERBB2* gene are found in several solid tumors, including breast cancer (Ahn et al., 2020). Several approaches have been developed to target HER2 (Cong et al., 2019). The monoclonal antibody (mAb) Trastuzumab was the first anti-HER2 agent approved in combination with conventional chemotherapy for patients with HER2-positive metastatic breast cancer and subsequently approved as adjuvant therapy in patients with early-stage disease (Boekhout et al., 2011). Trastuzumab binds to the HER2 extracellular domain, preventing receptor homo- or heterodimerization and blocking downstream activation. Taking advantage of HER2 as a model receptor, several EV functionalization strategies have been proposed, starting from the manipulation of EV-donor cells (*pre-isolation* strategies) or directly modifying EVs (*post-isolation* strategies). Different proteins have been exploited to anchor HER2-targeting moieties to EV surface, including transmembrane proteins or membrane-associated domains, also with Trastuzumab portions (Gurrieri and D’Agostino, 2022). However, the contribution of exposed receptors to the cell-targeting spectrum of vesicles remains elusive, especially of heterologous EVs interacting with different recipient cells. In this work, we exploited the CD81 protein, one of the most studied and structurally characterized tetraspanins (Kitadokoro et al., 2001), (Fordjour et al., 2022), to induce secretion of EV populations from HEK293T cells overexpressing CD81 fused with Trastuzumab light chains and a GFP reporter. After profiling the GFP-positive fractions in bulk EV populations and the retained binding capacity *in vitro* of the anti-HER2 moiety, we quantified EV-cell interactions using isogenic breast cancer cells with high or null HER2 expression, showing differences between isogenic cell lines manipulated for HER2 receptor expression. We functionally validated the EV-cell interaction by performing viability assays with secreted doxorubicin-EVs and direct detection of horizontally exchanged recombinant RNAs in mouse JIMT-1-engrafted tumors, demonstrating the need to decipher the multiple mechanisms underlying the interaction and consequent cargo delivery of heterologous EVs.

## Materials and methods

### Plasmids and cell lines

Trastuzumab light chains 1 and 2 were obtained by Tebubio Srl and cloned in the CD81-GFP vector (OriGene, 7268 bp), obtaining the antiHER2 construct (CD81-antiHER2-GFP, 7901 bp). Human embryonic kidney HEK293T (ATCC, CRL-3216), human breast cancer MDA-MB-231 (ATCC, HTB-26) and SK-BR-3 (AMSBIO, Abingdon, UK) cell lines were cultured under standard conditions in DMEM supplemented with 10% Fetal Bovine Serum, 2 mM L-Glutamine, and 100 U/ml penicillin-streptomycin (all Gibco). SK-BR-3 HER2-knockout (SK-BR-3 KO) were obtained using pSpCas9 BB-2A-Puro (PX459) V2.0 (9200 bp, Addgene) containing a sgRNA sequence (5’-TCATCGCTCACAACCAAGTG-3’) targeting exon 7 of *ERBB2* (cloned by Twin Helix) and selected with puromycin (Sigma-Aldrich) for 4 days (1 μg/ml puromycin for 72 hours, followed by 24 hours at 2 μg/ml). MDA-MB-231 cells expressing HER2 (MDA-MB-231 HER2+) were obtained by transient transfection of the pCMV3-SP-N-HA vector (6086 bp, SinoBiological) for 24 hours. Cells were transfected using either Lipofectamine 3000 according to manufacturer’s protocol (Invitrogen).

### Cell fractionation and immunoblotting

Cell fractionation experiments were performed as already described (Baghirova et al., 2015) in lysis buffer (50 mM HEPES pH 8, 10 mM NaCl, 10 mM MgCl_2_, 1 mM DTT, 10% glycerol, 1X protease inhibitor cocktail) supplemented by 25 µg/ml Digitonin (buffer A), 1% Igepal (buffer B) or 1% Triton X-100 and 1% Sodium deoxycholate (buffer C) for the sequential incubation and centrifugation protocol. Input samples corresponded to 2% of whole cell lysate from all the conditions analyzed. The first supernatant (cytosolic fraction) was collected after incubation of cells with buffer A on a rotary shaker for 10 min at 4 °C, then centrifuged at 2,000 rcf for 10 min at 4 °C. The obtained pellet was resuspended in ice-cold buffer B and vortexed before incubation on ice for 30 min and centrifuged at 7,000 rcf for 10 min at 4 °C. The resulting supernatant corresponded to the organelle-enriched fraction, while the pellet was resuspended in ice-cold buffer C with the addition of benzonase (Novagen) and incubated on a rotary shaker for 30 min at 4 °C. Next, samples were sonicated at 4 °C for 45 sec at 35 Amplitude (three cycles of 10 sec on and 5 sec off) within a ultrasonic bath sonicator (Q700, QSonica) and centrifuged at 7,800 rcf for 10 min at 4 °C to collect the nuclear fraction. Each fraction was loaded on 13% polyacrylamide gel for SDS-PAGE. Protein concentration was measured in triplicate with the bicinchoninic acid (BCA) protein assay kit (Thermo Fisher Scientific). Immunoblotting experiments were performed as already described(Notarangelo et al., 2019a) using the following antibodies: tGFP (TA150041, OriGene), SYNTENIN (ab133267, Abcam), CD9 (ab236630, Abcam), CALNEXIN (ab22595, Abcam), HER2 (ab237715, Abcam), RAB5 (C8B1, 3547, Cell Signaling Technology), GAPDH (GTX627408, GeneTex), H3 (GTX122148, GeneTex), SERCA2 (ab2861, Abcam), TSG101 (GTX70255, GeneTex), secondary antibodies Peroxidase AffiniPure Goat Anti-Mouse and Anti-Rabbit IgG (H+L) (Jackson ImmunoResearch).

### EV isolation and characterization

For EV collection, HEK293T were plated in 100 mm TC-treated Culture Dish (Corning) and transfected with 3 µg plasmid/dish when at 75% of confluence. Forty-eight hours later, cells were washed with PBS and incubated for additional 24 hours with serum-free medium. Media were centrifuged at 2,800 rcf for 15 min to eliminate cell debris and bigger particles. The supernatant was transferred to ultracentrifuge tube (38.5 ml-Beckman Coulter Ultra-Clear™) for ultracentrifugation at 100,000 rcf for 70 min at 4 °C using a Optima XE-90 ultracentrifuge with a SW 32 Ti rotor (Beckman Coulter). The EV pellet was resuspended in 0.22 μm-filtered sterile PBS and freshly-used or stored at −80 °C until further use. Nanoparticle tracking analysis (NTA) was performed using a NanoSight NS300 instrument (Malvern Panalytical, Malvern, UK) with a 532 nm laser. Each sample was subjected to 3-5 consecutive 60 sec videos recorded at camera level 15. Particle concentration and size distribution were determined using NanoSight NS300 software NTA 3.4 Build 3.4.003 (Malvern Panalytical), setting 4 as a detection threshold.

Imaging flow cytometry was performed using an ImageStreamx (ISX) MKII instrument (Luminex Corporation) at 60x magnification, high gain mode, and low flow rate. EV samples in PBS were labeled with 1 μg/ml (final concentration) of Cell Mask Deep Red (CMDR, Invitrogen) in ratio 1:1 (v/v) and incubated at RT for 20 min. Then, samples were diluted in PBS to obtain a final concentration lower than 10^10 objects/ml before acquisition. All samples were analyzed using INSPIRE® software (Luminex Corporation, Seattle, WA, USA), with a minimum of 3,000 events collected. Data analyses were performed using ISx Data Exploration and Analysis Software (IDEAS®, Luminex Corporation, Seattle, WA, USA). Cryogenic electron microscopy (Cryo-EM) acquisitions were performed at Fisher Scientific (Eindhoven, The Netherlands). QuantiFoil 1.2/1.3 cu 200 grids, pretreated with 30 sec glow discharging, were used. Vitrobot parameters were set as follows: 3.0 μl of each sample, temperature 4 °C, humidity 95%, blot time 7 sec, blot force 0. Fifteen cryo-electron images were collected per each sample using a transmission electron microscope Glacios (Thermo Fisher Scientific), equipped with a Falcon 4i Selectris camera, at a nominal magnification of 49,000x. Imaging parameters were set as follows: Pixel size (Å) 2.4, Dose rate (e/pix/sec) 12.4, Total dose (e/Å^2) 20, Exposure time (sec) 9.3, Energy filter (eV) 10, defocus −1.9 μm. For image analysis, vesicular structures were manually selected as regions of interest (ROIs) using FIJI software and the size measured in pixels. From the ROI’s pixel size, the actual area and diameter of EVs were calculated.

### AlphaLISA interaction assay

The Amplified Luminescent Proximity Homogeneous Assay (ALPHA Assay) was performed in white 384-well Optiplates (PerkinElmer) in a final volume of 20 μl. The antiHER2 protein was produced from CD81-antiHER2-tGFP vector (600 ng) using TNT® Quick Coupled Transcription/Translation Systems (Promega), following manufacturer’s instructions. The resulting product was pre-incubated with 30 nM HER2-DDK (TP322909, OriGene) for 30 min before addition to the other components: 10 nM tGFP Ab (TA150041, OriGene), anti-FLAG donor and protein G coated acceptor beads (10 ng/μl final concentration; AS103D and AL102C, PerkinElmer). For the competitive assay with CD81-GFP and antiHER2 EVs, 40 nM HER2-DDK and 9 nM anti-HER2 Ab (ab237715, Abcam) were used. All the components were diluted in the Alpha Screen Control Buffer (Perkin Elmer). The signal was detected using an EnSight® multimode plate reader (PerkinElmer) after 1 hour of plate incubation at RT in the dark under rotation at 70 rpm.

### Immunoprecipitation and immunofluorescence

Serum-free media from transfected HEK293T was concentrated to about 2 ml with Amicon Ultra-15 Centrifugal Filter 10 kDa MWCO (Merck Millipore) at 3,000 rcf for 15-20 min, and diluted according to the relative concentration of GFP-positive particles. Input samples were prepared from 15 μl of undiluted conditioned medium. Anti-FLAG M2 magnetic beads (Sigma-Aldrich) or protein G dynabeads (Invitrogen) were washed three times in PBS before pre-incubation, at 4 °C for 1.5 hours under rotation, with HER2-DDK or tGFP antibody, respectively. Bead-antibody mixtures were added to the media for 20-30 min at 4 °C under gentle rotation. For competitive HER2 binding, 1 μg/sample of Trastuzumab (anti-HER2-Tra-hIgG1, InvivoGen) was added directly to the media before incubation with beads. After the final incubation, beads were washed with PBS and resuspended with 1X Laemmli sample buffer for subsequent denaturation at 98°C for 5 min and SDS-PAGE.

Immunofluorescence for HER2 detection was performed in breast cancer cell lines fixed with 4% paraformaldehyde for 15=min at RT and washed three times with cold PBS. Blocking (10% FBS, 0.05% Triton X-100 in PBS) was performed for 1 hour at RT followed by primary Ab (ab237715, Abcam, diluted 1:1000, 0.1% FBS in PBS) incubation for 1 hour at RT. After three washes with PBS, cells were incubated for 1 hour at RT with Goat anti-Rabbit Alexa Fluor™ 633 (Invitrogen, diluted 1:1000, 0.1% FBS in PBS). Three washes with cold PBS 5 min each were performed before the addition of Hoechst for 15 min. One final wash with cold PBS was performed before acquisition. For CD81 and RAB5 IF, HEK293T were seeded on optical coverslips and IF was similarly performed, with the addition of a 5 min permeabilization step (0.1% Triton X-100 in PBS) before primary Ab incubation overnight at 4=°C (0.1% FBS and 0.05% Triton X-100 in PBS). CD81 Ab (MA5-13548, Invitrogen) or RAB5 Ab (C8B1, 3547, Cell Signaling Technology) were used as primary Ab, Goat anti-Mouse or anti-Rabbit IgG (H+L) Cross-Adsorbed Secondary Antibody Alexa Fluor™ 568 (Invitrogen). Coverslips were mounted using ProLong Diamond Antifade Mountant (Invitrogen).

### Confocal microscopy and image analysis

Time-lapses and images of EV uptake by recipient breast cancer cells were acquired at the Optical Imaging Centre (OIC) at Erasmus MC (Rotterdam, The Netherlands) with a LEICA TCS SP8 AOBS confocal microscope, with Galvo Z stage and Adaptive Focus Control, using a HC Plan Apo CS2 40x/1.3 oil immersion objective. Live cell confocal imaging was performed under humidified conditions with 5% CO2 at 37 °C. MDA-MB-231 and SK-BR-3 cells were seeded in glass-bottom dishes (CELLview™ Culture dish, 35 mm, four chambers) and, before acquisition, 340 μl DMEM already containing 1.25 nM LysoTracker™ Red DND-99 (L7528, Invitrogen) and 1 µg/ml Hoechst (62249, Thermo Scientific) were added to each chamber for 15 min. Before starting acquisition CD81-GFP or antiHER2 EVs were added in a ratio of 20,000-50,000 per seeded cell, in 350 μl as final volume. For fixed cell acquisitions, MDA-MB-231 (wt and HER2+) and SK-BR-3 (wt and KO) were seeded in the same dishes and incubated with EVs for 4 hours considering the relative abundance of GFP-positive EVs, then washed with PBS before fixation and immunofluorescence (IF) for HER2 receptor. Fourteen Z-stacks were acquired within around 11 μm of total Z size, with voxel size 0.1623×0.1623×0.7991 μm3. Images were processed and analyzed using FIJI/ImageJ software.

To quantify EV-cell interactions, an automated pipeline was applied using CellProfilerTM version 4.0.7 on the Maximum Intensity Projection of 14 z-stacks for each acquired channel. Cell nuclei were identified as primary objects with a threshold in the Hoechst blue channel, the cytoplasms were defined as secondary objects using a low threshold in the HER2 red channel that allowed the segmentation of a cytoplasmic region also in the HER2 negative cells. EVs were defined as green spots in a range of diameters from 0.8 to 5 µm and the interaction within the cells has been established overlapping the EVs object mask to the cytoplasmic region using the object processing function “RelateObject”. Objects number, AreaShape and Red Intensity features were calculated for the identified Nuclei, Cytoplasms and EVs. The final Spreadsheets have been combined and the dataset has been analyzed using KNIME Analytics Platform v 4.7.5.

Images in Figure 1A-B were acquired with a Nikon AX laser scanning inverted confocal microscope, using a Plan Apo H 60x/1.4 oil immersion objective. Images in Figure 5A and Supplementary Figure 3C were acquired with a Nikon Ti2 inverted microscope equipped with a Crest X-light V2 Spinning Disc system and an Andor iXon Ultra 888 EMCCD camera, using a Plan Apo 20x/0.75 objective. Images in Supplementary Figure 1B were acquired with a Nikon Ti2 inverted microscope equipped with a Crest X-light V2 Spinning Disc system and an Andor Zyla 4.2 PLUS sCMOS camera, using a Plan Apo 20x/0.75 objective. For all the acquisitions, settings were kept constant within the same experiment and linear adjustments for brightness and contrast were equally applied to the reported images.

**Figure 1:**
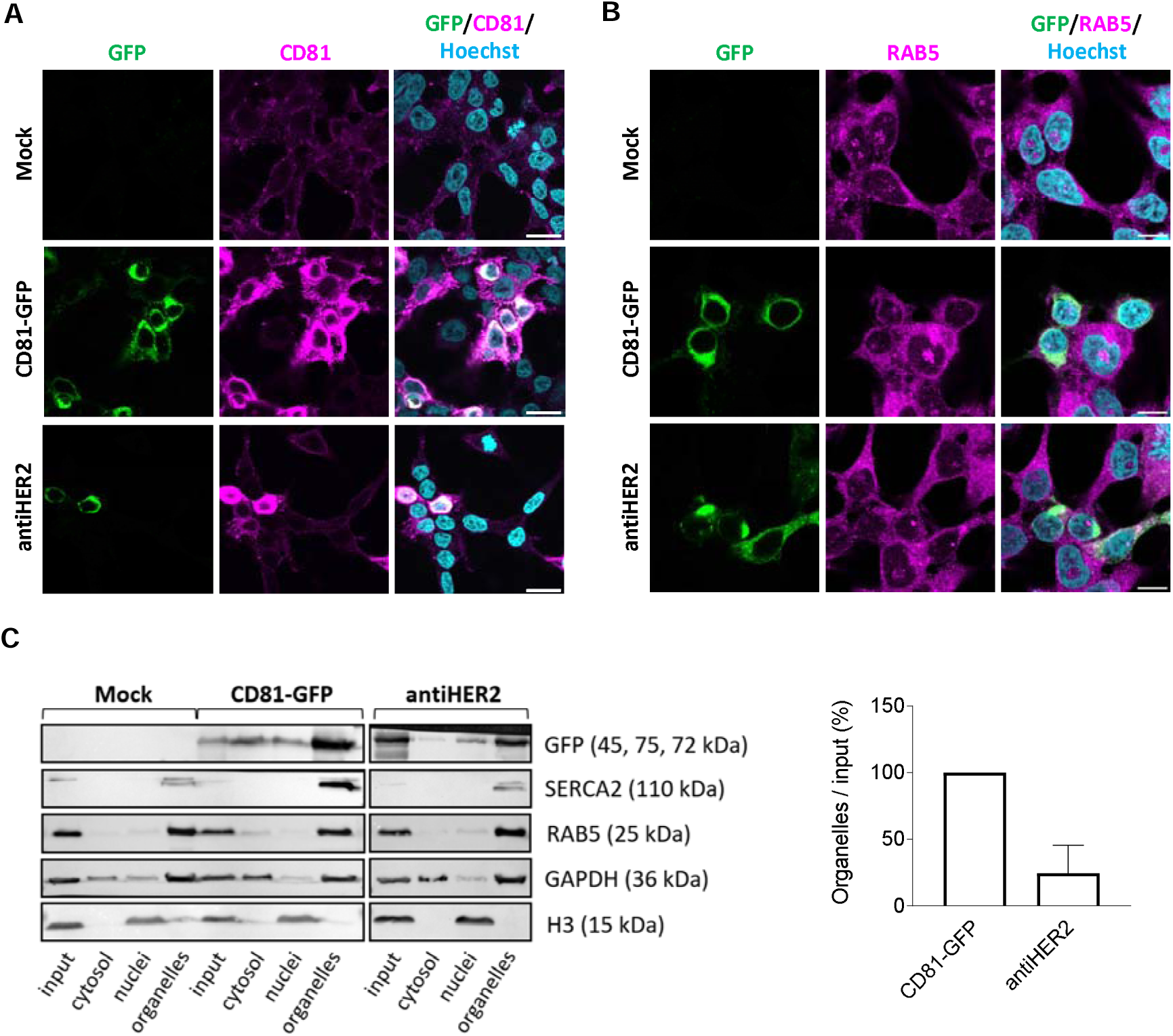
CD81 fusion proteins are expressed in HEK293T upon transient transfection and co-sediment with organelle-enriched sub-cellular fractions. **A-B)** GFP detection and immunofluorescence staining of endogenous CD81 and RAB5 proteins in transfected HEK293T cells. Cell were subjected to confocal microscopy after 48 hr of transfection with CD81-GFP and antiHER2 plasmids. Recombinant proteins are visualized in green (GFP), endogenous CD81 or RAB5 in magenta (Alexa Fluor 568), and cell nuclei in cyan (Hoechst). Scale bar is 20 μm in A and 10 μm in B. **C)** Immunoblotting of sub-cellular fractions obtained through a sequential lysis buffer-centrifugation protocol. Separation of subcellular fractions was confirmed by the enrichment of corresponding protein markers: Cytosol (GAPDH), nuclei (histone H3), and organelles (SERCA2 for endoplasmic reticulum, RAB5 for early-endosomes). GFP-positive chimeric proteins were detected at the expected molecular weight (45 for CD81-GFP and 75 kDa for antiHER2). The histogram reports the densitometric quantification with mean and SD of two independent experiments.

### RNA isolation and digital droplet PCR

Total EV-RNA was extracted with TRIzol reagent (Ambion, Life Technologies) and chloroform precipitation, followed by single-cell RNA purification (Norgen kit) including on-column DNAse I (Qiagen) treatment for 10 min at RT. cDNA synthesis was performed following manufacturer’s instructions (SensiFAST™, Meridian Bioscience™) starting from 15 μl of RNA template. To assess the integrity of the fusion protein-encoding transcripts (CD81-GFP, antiHER2), the corresponding cDNA samples were amplified (Supplementary Table 2) prior DNA electrophoresis. ddPCR experiments were carried out using EvaGreen, following manufacturer’s instructions (Bio-Rad). EV-derived cDNA samples (5.5 μl each) were mixed with 11 μl of 2X QX200™ ddPCR™ EvaGreen Supermix (Bio-Rad) and 5.5 μl of primers (35 nM each). The following primers were used to specifically amplify the fusion sequences of interest: A) 5’-CTTCAAGGAGGACTGCCAC & 5’-GCTCGCGCTATAAATCAGCAGT; B) 5’-TGACCAAAAGCTTTAACCGTG & 5’-TGGGGTAGGTGCCGAAGT; C) 5’-CTTCAAGGAGGACTGCCAC & and 5’-TGGGGTAGGTGCCGAAGT) and target concentration was determined using QuantaSoft Software™ (Bio-Rad).

RNA extraction from tumor xenografts was performed with TRIzol and RNeasy Mini Kit (Qiagen), following the manufacturer’s instructions. cDNA synthesis was carried out using WarmStart reverse transcriptase (NEB, M03805) from 6 μg of RNA as starting material, including couple B primers (160 nM each). RNAse H from E. coli (Illumina) was added (10U/sample) for 30 min at 37 °C. cDNA purification was performed using the Nucleospin gel and PCR clean-up kit (Macherey-Nagel, 740609.50) following manufacturer’s instructions. ddPCR reactions were performed with cDNA diluted 1:300.

### UHPLC-MS analysis of secreted doxorubicin

To collect doxo-EVs from drug-treated cells, transfected HEK293T were incubated with 10 μM doxorubicin (BD32885, BLD Pharmatech GmbH) for 3 hours at 37 °C and 5% CO2. After a PBS wash, EVs were isolated from serum-free DMEM as previously described. Metabolite extraction was carried out by adding 80% cold methanol to the EV stocks. Samples were then vigorously shaken (5 min) and kept at −80°C overnight. Finally, the samples were vacuum-dried using a SpeedVac concentrator. Dried extracts were equilibrated to RT, resuspended in 30 µl of acetonitrile:water:formic acid (5:95:0.1%, v/v/v), and thoroughly mixed. Seven serial drug dilutions, ranging from 8,000 to 1.95 nM, were used to prepare the standards for the calibration curves. Ten μl of standards and samples were injected onto Ultimate 3000RS (Thermo Scientific) UHPLC system coupled online with an Orbitrap Fusion Tribrid (Thermo Scientific) mass spectrometer. A Hypersil Gold C18 column (Thermofisher, 100 x 2.1 mm, particle size: 1.9 µm) was used for separation. The LC method consisted of a linear gradient from 5 to 100% B (B: acetonitrile 0.1% of formic acid; A: water + 0.1% formic acid) over 15 min, followed by 4 min at 100% at the flow of 0.2 ml/min. The MS spray voltage was set at +3500V with the ion transfer tube temperature set at 300 °C (sheath and auxiliary gasses were set at 20 and 5 Arb, respectively). The MS data were acquired in full scan in the Orbitrap at 120.000 FWHM (200 m/z), in the scan range of 100-1000 m/z. Software FreeStyle ver.1.6 (Thermo Scientific) was used to inspect mass spectra. The area under the peak of each precursor ion and the total ion current (TIC) were extracted using Skyline (MacCoss Lab Software)(Adams et al., 2020). The software derived precise m/z values as well as isotope distributions for each precursor. Each analyte was also investigated for common adducts, [M+H]+, [M+K]+, [M+NH4]+, [M+Na]+, and the [M−H2O+H]+ ions, for each considering the three most abundant isotopes. The total area of precursor ions was calculated by summing all precursor levels. The log2-transformed value of the total area of precursor ions of each standard sample was plotted as function of the concentration to construct the standard curves of each compound.

### Cell viability assay

The 3-(4,5-dimethylthiazol-2-yl)-2,5-diphenyl-2H-tetrazolium bromide (MTT, Thermo Scientific) assay was performed on cells seeded and treated in 96 plates for 72 hours with the free doxorubicin or doxo-EVs. Doxorubicin concentration in doxo-EVs was estimated from previous LC-MS analysis. After incubation, the medium was removed and MTT solution (0.5 mg/ml in DMEM) incubated for 4 hours at 37 °C and 5% CO2, before cell lysis in DMSO. Absorbance (570 nm) was measured at a Varioskan LUX Multimode Microplate Reader (Thermo Fisher Scientific) and cell viability was calculated as % with respect to untreated cells (DMEM only).

### In vivo study

The *in vivo* study was performed in collaboration with Reaction Biology Europe GmbH (Freiburg, Germany). Each experimental groups contained five female athymic nude mice (Crl:NU(NCr)-Foxn1nu). On Day 0, 5.0 × 106 JIMT-1 human breast carcinoma tumor cells in 100 µl PBS were implanted into the left mammary fat pad of each mouse. After animals had been randomized on Day 12, treatments of the test samples were initiated. All treatments were administered at a dose of 0.5 µg/kg and a dosing volume of 5 ml/kg subcutaneously at the tumor implantation site. Animal weights were measured three times, one time on day of randomization (Day 12) and two times after the start of therapy (Days 13 and 15). During the study, the growth of the intramammary implanted JIMT-1 primary tumors was determined twice by caliper measurement on Days 12 and 15. All animals reached the end of the study as scheduled. The primary tumor samples were collected during the final necropsy on Day 15, the study endpoint.

### Statistical analysis

Statistical analysis was performed using GraphPad Prism 9, as well as data visualization, applying non-parametric Anova Kruskal-Wallis or Student’s t-test. A minimum of 95% confidence level was considered significant. The significance level was set at *: p < 0,05; **: p <0.01; ***: p<0.001; ns: not statistically significant. Mean and standard deviation of independent experiments are reported in the graphs and detailed in figure legends. Representations were created in biorender.com, where indicated.

## Results

### CD81 fusion proteins participate in intracellular vesicular trafficking

Recombinant plasmids were designed to encode a human full-length CD81 protein in frame with the turbo GFP reporter (CD81-GFP) and including Trastuzumab light chains 1 and 2 (CD81-antiHER2-GFP, or antiHER2), generating two fusion proteins with an expected molecular weight of 45 and 75 kDa, respectively (**Supplementary Figure 1A**). After 48 hours of HEK293T cell transfection, the CD81-GFP protein showed the highest fluorescence intensity, which was slightly reduced in the case of antiHER2. In this setting, chimeric proteins populated the perinuclear region, including a partial granular cytoplasmic and plasma membrane distribution matching the endogenous CD81 protein, in contrast to the whole cell body stained by GFP alone (**Figure 1A and Supplementary Figure 1B**). By counterstaining cells with an antibody recognizing RAB5, a marker of early endosomes (Shearer & Petersen, 2019; Mathieu et al., 2021), we found circumstantial spots overlapping with GFP in confocal microscopy (**Figure 1B**). Subsequently, cell fractionation and immunoblotting experiments (**Figure 1C**) indicated a co-sedimentation of fusion proteins in RAB5/Sarco-Endoplasmic Reticulum Calcium ATPase (SERCA)-positive (organelle-enriched) fractions, supporting the potential involvement of tetraspanin-guided fusion proteins in vesicular trafficking.

### CD81 fusion proteins are detected on secreted EV populations

To investigate the presence of chimeric proteins in cell-secreted EVs, transfected HEK293T cells were exposed to serum-free media for 24 hours. Media were subjected to differential ultracentrifugation (DUC), and recovered particles were characterized by nanoparticle tracking analysis (NTA), cryogenic electron microscopy (Cryo-EM), immunoblotting, and imaging flow cytometry. NTA experiments showed a heterogeneous particle size distribution in all the conditions analyzed, with the most represented populations between 100 and 200 nm (**Figure 2A**). Although no significant changes were detected in mode diameters, the mean diameters showed a rising trend with ectopic CD81 proteins, contributing to a significant shift of about 15-25 nm in the case of antiHER2 particles (**Figure 2B**). In parallel, the two ectopic proteins, especially antiHER2 (p value=0.0005), were associated with a significant increase in particle concentration compared with Mock samples (**Figure 2B**).

**Figure 2:**
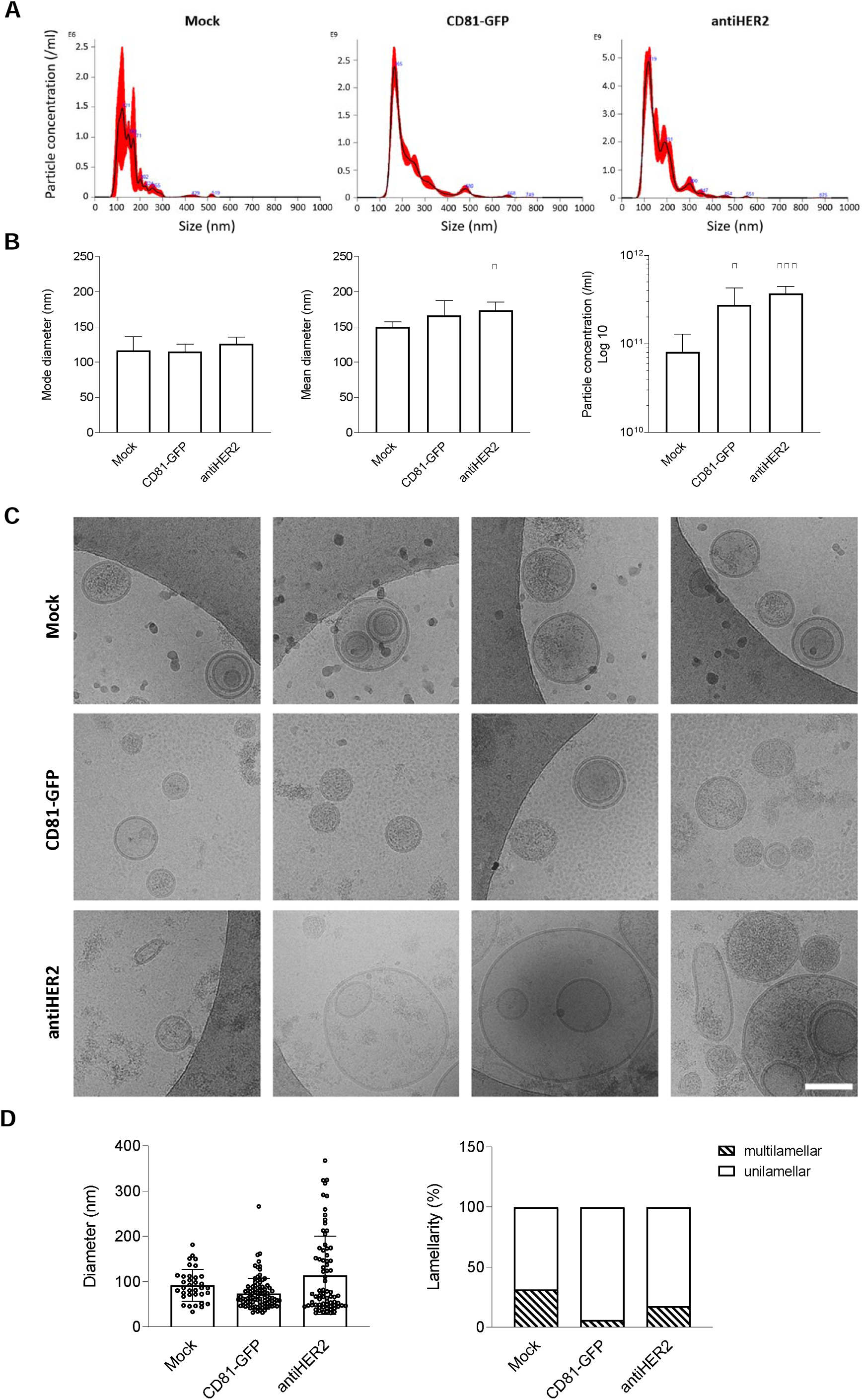
Profiling of particles recovered from transfected HEK293T cells. **A-B)** Nanoparticle Tracking Analysis (NTA) of particles secreted by transfected HEK293T cells. Representative size distribution profiles of Mock, CD81-GFP, anti-HER2 samples. The black curve indicates the mean of three measurements, with SE in red. Mode and Mean diameters, and particle concentration are plotted. Error bars include at least three biological replicates. Significance * is P<0.05, *** P<0.001, vs Mock condition. **C)** Representative Cryo-EM images of Mock, CD81-GFP and antiHER2 EV samples confirming the vesicular structure and size heterogeneity of recovered vesicles. The indicated scale bar is 100 nm. **D)** Plot of the observed diameter of vesicles in Cryo-EM images (n=35 for Mock, n=99 for CD81-GFP, n=74 for antiHER2) and lamellarity, expressed as percentage of unilamellar and multilamellar vesicles over the observed bulk EV populations.

To gain insights into the quality of released EVs, we acquired Cryo-EM images (**Figure 2C**). Morphological analyses indicated an almost intact lipid bilayer with a predominant spherical shape for most retrieved particles. Notably, the imaging analysis revealed subsets of larger vesicles in the antiHER2 condition (**Figure 2D**), mirroring the results obtained by NTA. In addition, Cryo-EM acquisitions systematically showed multi-layered (or multi-lamellar) vesicles, representing 10-30% of total EVs, with a trend of decrease in the ectopic conditions compared to Mock (**Figure 2D**). Immunoblots on the same particle samples revealed the presence of chimeric proteins at the corresponding molecular weight, with concomitant accumulation of CD9 and SYNTENIN EV-markers, in contrast to EV-negative marker CALNEXIN, which was undetected when compared to cell lysates (**Figure 3A**). To further substantiate fusion proteins as part of EV protein cargo, we analyzed non-denatured samples by imaging flow cytometry using Cell Mask Deep Red (CMDR), a fluorescently-labeled dye with an affinity for lipid membranes. Once gated for CMDR positivity, the fraction of GFP-positive EVs was quantified (**Figure 3B**). CD81-GFP EVs presented the highest percentage of double-positive events compared to antiHER2 EVs (66.5±4.7% and 15.65±8.76%), with a consistent no signal in the Mock vesicles. Overall, CD81 fusion proteins were confirmed as cargoes of heterogeneous EV populations.

**Figure 3.**
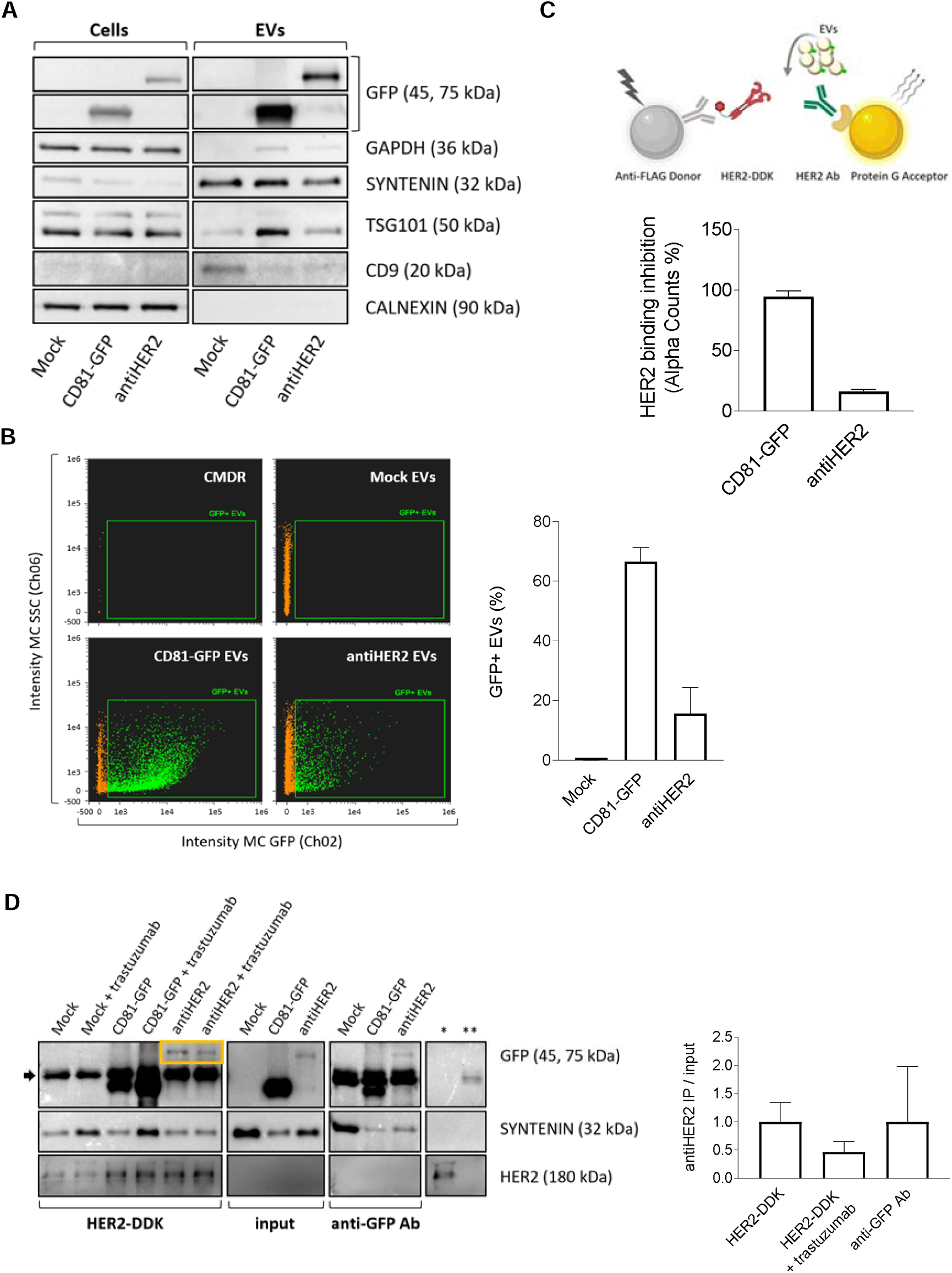
CD81-guided fusion proteins are cargo of secreted EV populations. **A)** Representative immunoblotting of cell and EV lysates (1 μg proteins/well). EVs are positive to transmembrane (CD9) and cytosolic proteins (SYNTENIN, TSG101), while negative to CALNEXIN, and with low detectable levels of GAPDH compared to cell lysates. **B)**. Dot plots of imaging flow cytometry to detect GFP-positive EVs. The green fluorescent signal (Ch 2, 488 nm laser) was detected as sub-gating of EVs labeled with Cell Mask Deep Red (CMDR, in orange, Ch 11, 635 nm) to side-scatter (Ch 6). Non-fluorescent, calibrator SpeedBeads, Amnis (1 µm) were continuously run during acquisitions. The graph shows the quantification of double-positive particles. Mean and error bars derive from three independent experiments. **C)** Sandwich designed for the AlphaLISA competitive assay. CD81-GFP and antiHER2 EVs were tested for competition with HER2-DDK. Image created with BioRender.com. The graph shows the measured alpha counts normalized to the GFP-positive EV population as calculated by NTA and imaging flow cytometry. Mean and SD derive from three independent experiments (significance is **** P<0.0001). **D)** Representative western blotting of recombinant EVs immunoprecipitation with HER2-DDK or anti-GFP antibody in serum-free DMEM. GFP-positive fusion proteins are enclosed in the yellow box above the antibody heavy chains (black arrow), SYNTENIN, and HER2-DDK. Controls of beads flow through with HER2-DDK (*) or anti-GFP antibody (**) are shown on the right, indicating saturation of the beads’ surface to avoid non-specific binding. The graph shows the densitometric quantification of antiHER2 EVs captured by both HER2-DDK and anti-GFP Ab, with a competition effect of Trastuzumab. Mean and SD refer to two independent experiments.

### AntiHER2 fusion protein retains a HER2 receptor binding capacity *in vitro*

Since the mature endogenous CD81 protein is engaged in vesicular membranes(Fan et al., 2023), we functionally investigated the exposure of CD81 fusion proteins on recovered EV populations. To this aim, we tested the HER2-receptor binding capacity of secreted EVs with fusion proteins retaining the GFP reporter. We developed an AlphaLISA assay using a human DDK (or FLAG)-tagged HER2 recognized by anti-FLAG Donor beads and an *in vitro*-translated antiHER2 protein recognized by an anti-GFP antibody coupled with Protein G Acceptor beads (**Supplementary Figure 2A**). We detected a specific ligand binding that was further improved by shortening the interacting sandwich using an anti-HER2 antibody (**Supplementary Figure 2B**). With these ligands, we optimized the bead:ligand ratio (hooking point) to perform competitive assays based on the interference of secreted antiHER2 EVs compared with CD81-GFP EVs. The antiHER2 EVs remarkably reduced the specific antibody:HER2 interaction (p value<0.0001), confirming a fraction of secreted EVs indeed exposing a functional moiety that, despite the presence of GFP, retained a measurable binding activity and competed with a canonical anti-HER2 antibody (**Figure 3C**). Interestingly, we obtained the same indication from immunoprecipitation (IP) experiments using HER2-DDK or anti-GFP antibody to capture the fusion proteins on EVs. HER2-DDK/anti-FLAG or anti-GFP/Protein G beads were incubated with EV-containing media diluted according to the relative particle concentration detected by ImageStream. IP-GFP bands at the corresponding size and SYNTENIN bands confirmed the presence of secreted EVs exposing the recombinant proteins with accessible GFP and Trastuzumab light chains moieties (**Figure 3D**). The canonical Trastuzumab was also included in these experiments as a fourth ligand to investigate competition effects. Notably, antiHER2 EVs, but not CD81-GFP EVs, reproducibly competed with Trastuzumab, reducing the IP signal by about 50%. Although we cannot exclude a sub-optimal interaction due to antibody-mediated aggregation and the presence of GFP, antiHER2 EVs were capable of specific interactions with a human recombinant HER2 receptor. These results encouraged the use of these EV populations (from now on referred to as “recombinant EVs”) for EV-cell interaction studies.

### Recombinant EVs show heterogeneous interactions with breast cancer cells

By confocal and live cell imaging, we studied the *in vitro* tropism of HEK293T-derived EVs on different breast cancer cells. First, we checked the selective receptor expression of HER2-positive SK-BR-3 and HER2-negative MDA-MB-231 cells (**Figure 4A and Supplementary Figure 3A**). Next, to better investigate the biological relevance of the receptor on potential EV-cell interactions, we ectopically expressed HER2 in MDA-MB-231 to obtain HER2-positive cell populations. On the other hand, we selected CRISPR/Cas9-mediated *ERBB2-*KO SK-BR-3 cells to obtain clones with an abrogated receptor (**Figure 4B and Supplementary Figure 3B**). In parental cells, recombinant EVs were tracked up to ten hours of incubation with living cells grown under standard conditions. A ratio of about 30,000 bulk EVs per seeded cell was maintained to ensure GFP detection (**Supplementary Figure 2D**). Time-lapse confocal imaging (**Figure 4C**) showed EV-cell interaction and internalization followed by partial lysosome accumulation, as demonstrated by some LysoTracker co-localizing spots. These results confirm that heterologous EVs could undergo different fates upon endocytosis, including lysosomal degradation or recycling(Herrmann et al., 2021).

**Figure 4:**
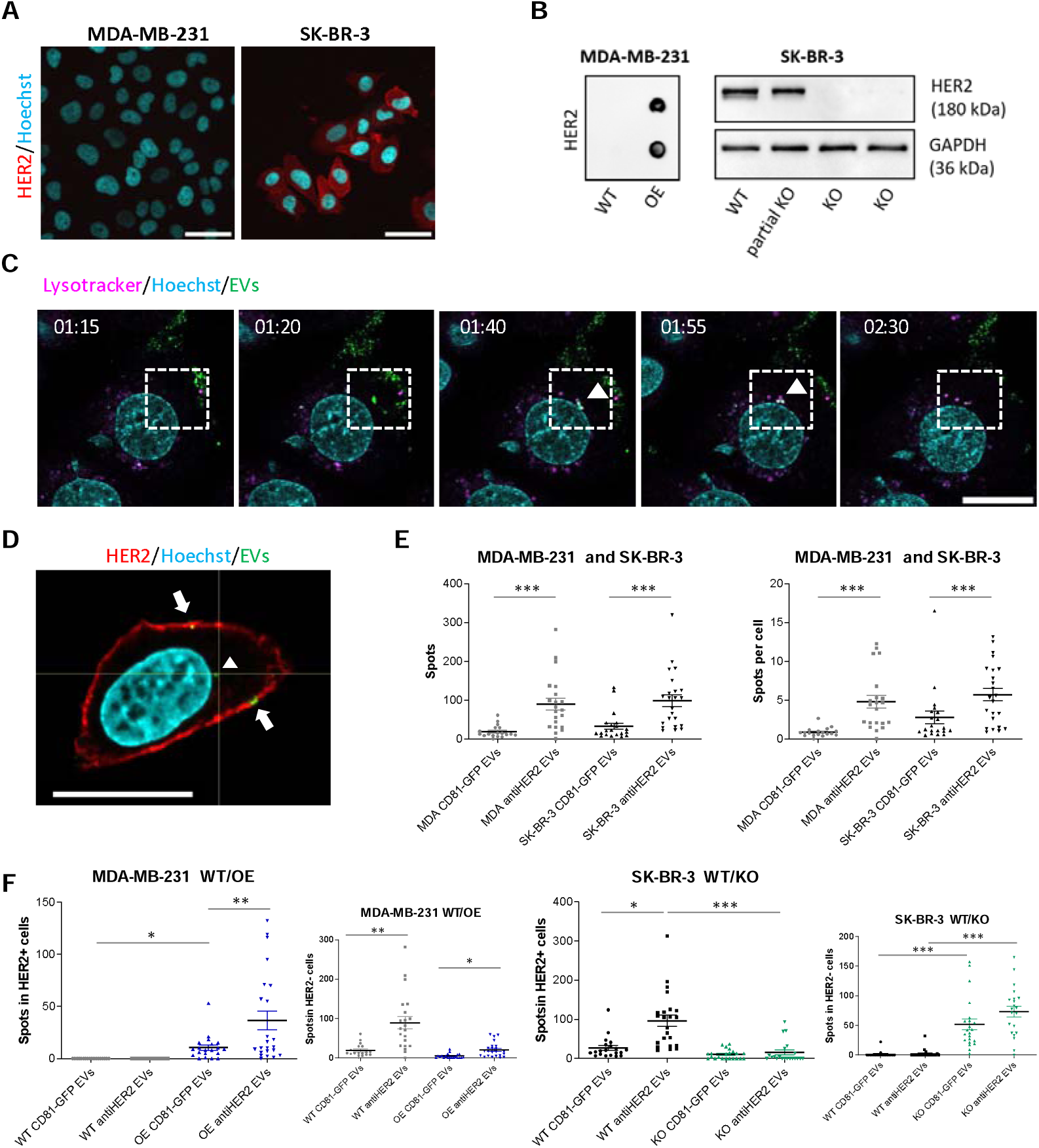
Recombinant EVs show heterogeneous interactions with breast cancer cells. **A)** Immunofluorescence of MDA-MB-231 and SK-BR-3 breast cancer cell lines as HER2 negative or positive cells, respectively. HER2 receptor is in red (Alexa Fluor 633), nuclei are shown in cyan (Hoechst). The indicated scale bar is 50 μm. **B)** Left: HER2 protein detection by Dot blot in lysates from wild-type or transfected (OE) MDA-MB-231 cells. Right: Immunoblot for checking the selection of SK-BR-3 cells with HER abrogation (*ERBB2*-KO, or KO). **C)** Representative confocal time lapse of recombinant EVs incubated with live cells (time points are indicated). GFP-EVs are shown in green, lysosomes are shown in magenta (Lysotracker red), and nuclei in cyan (Hoechst). The white squares highlight the co-localization between EVs and lysosomes (white arrowhead). The indicated scale bar is 20 μm. **D)** Representative confocal image of fixed SK-BR-3 cells recognizing HER2 (Alexa Fluor 633) and nuclei (Hoechst) after 4 hr incubation with CD81-GFP EVs (green spots). White arrows or the arrowhead indicate different localization of EVs upon cell interaction. Indicated scale bar is 20 μm. **E-F)** Quantification of recombinant EVs with recipient breast cancer cell lines. MDA-MB-231 (WT and HER2 OE) and SK-BR-3 (WT and KO) were incubated with EVs for 4 hours, then washed with PBS before fixation and HER2 immunofluorescence. Fourteen Z-stacks were acquired within around 11 μm of total Z-size and the Maximum Intensity Projections have been analyzed with an automated pipeline (using CellProfilerTM 4.0.7). Graphs report Mean and SD of the spot distribution from three independent experiments (* if P<0.05, ** if P<0.01, *** if P<0.001).

In parallel, cells were incubated with recombinant EVs for 4 hours before fixation, washing, and imaging. We performed a high-content imaging analysis on Z-stack maximum intensity projection by quantifying GFP spots in a single cell area defined by the HER2 receptor (**Figure 4D**). The exposure of both MDA-MB-231 and SK-BR-3 resulted in an equivalent cumulative number of GFP spots, although antiHER2 EVs appeared more prone to interact compared to CD81-GFP EVs (**Figure 4E**). Interestingly, this trend remained when normalizing the number of spots against the number of EV-receiving cells, indicating an EV-cell interactome substantially independent of the exposure of the HER2 receptor in these cells. However, the tropism of recombinant EVs was significantly influenced by receptor dosage in isogenic cells, dictating a positive distribution of GFP spots in HER2-positive MDA-MB-231 cells and a dramatically reduced signal in the SK-BR-3 cells with abrogated HER2 receptor (**Figure 4F**). The observed distribution was paralleled by complementary spot numbers detected in HER2-negative cell populations. These results suggest the existence of multiple mechanisms of EV-cell interactions with potential cell lineage-dependent dynamics. Therefore, the heterologous EV-homing and the presence of a specific surface receptor could partially contribute to the recruiting mechanisms that determine the EV tropism/biodistribution.

### HER2 receptor exposure influences the sensitivity to heterologous doxorubicin-EVs

To functionally investigate the consequences of heterologous EV-cell interactions, we applied an EV pre-isolation strategy to optimize the recovery of vesicles with co-secreted doxorubicin (doxo-EVs)(Sritharan and Sivalingam, 2021), (Farhat et al., 2022). The main steps we followed included doxorubicin treatment of transfected HEK293T cells with different doses and timing, particle sedimentation by DUC, and mass spectrometry to detect doxorubicin. We tested three drug concentrations (0.5, 5, and 15 µM) along with treatment (3 or 9 hours) and release-kinetic schedules (6, 20, or 24 hours, **Supplementary Figure 4A**). We then selected ten µM for 3 hours of treatment and 24 hours of release as an acceptable protocol for preserving cell viability, doxorubicin abundance, and NTA profiles. The relative concentration of doxorubicin per particle was determined through a calibration curve (**Supplementary Figure 4B**) run in parallel with NTA analyses (**Supplementary Figure 4C**). The number of recovered particles correlated with doxorubicin concentration, despite a variability we observed in the range of 0.3 to 4 µg per 10^12^ particles among the biological preparations (**Supplementary Figure 4D-E**). Given their stable growth after antibiotic selection and differential behavior with recombinant EVs, we performed viability assays with wild-type and HER2-KO SK-BR-3 cells. Treatment with free doxorubicin caused similar cytotoxic effects on these cells, with a calculated IC_50_ of 185 and 186 nM, respectively (**Figure 5A**). Then, we performed MTT assays by treating cells with secreted doxo-EVs for 72 hours with a calculated drug concentration of 60 nM. We included Mock-doxo EVs and free drugs (doxorubicin and DMSO) as controls (**Figure 5B**). In settings where the free drug only reduced the cell viability by 10%, doxo-EVs killed almost 50-65% of wild-type cells, demonstrating an increased bioactivity of the co-secreted drug. In line with the observed EV-cell interactome, antiHER2-doxo EVs showed slightly better efficacy than CD81-GFP-doxo EVs but were far less effective in HER2 KO cells. These data indicate that exposed receptors can functionally influence the interaction spectrum of heterologous EVs with subsequent cargo internalization/bioavailability.

**Figure 5:**
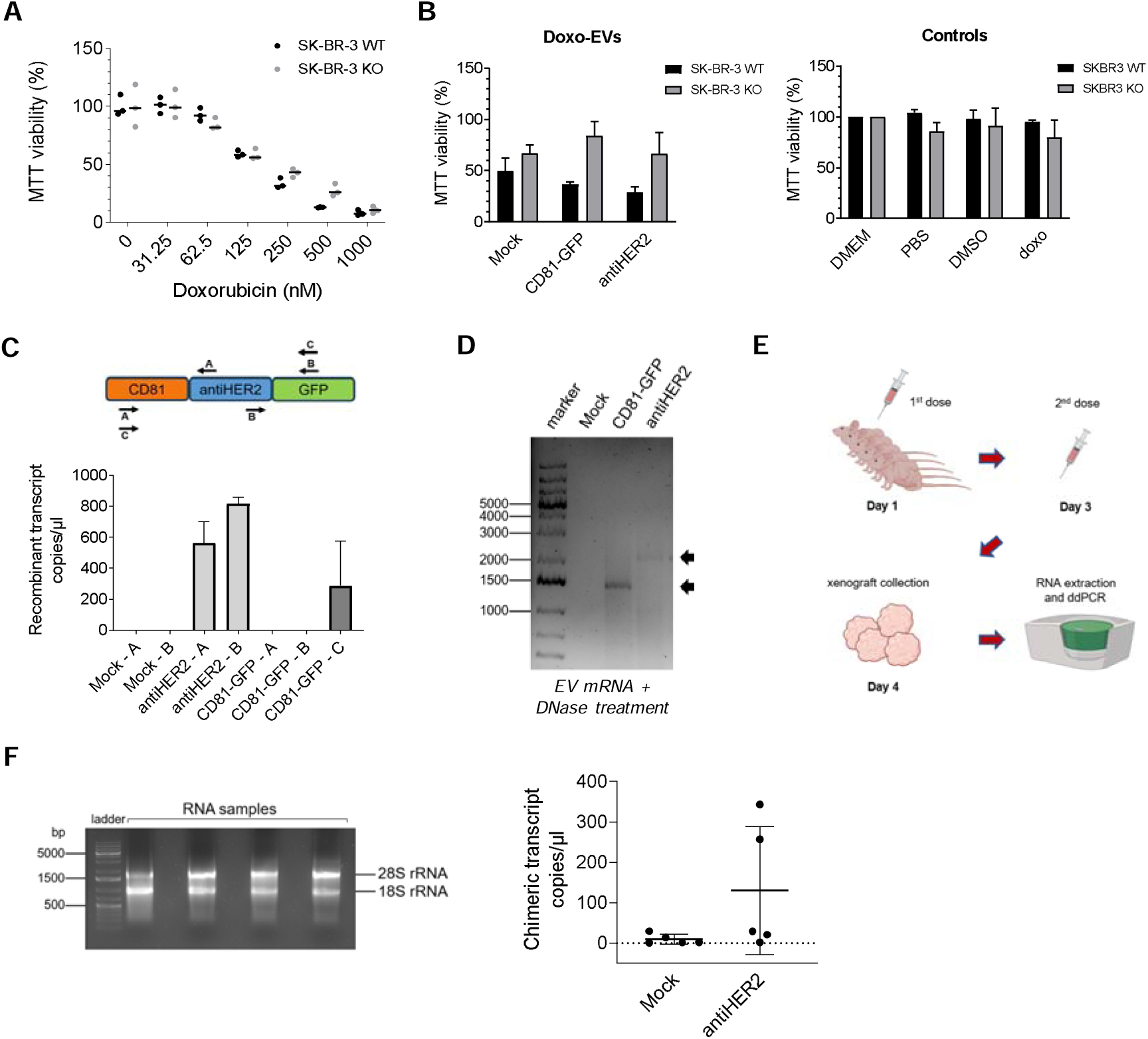
Functional consequences of recombinant EV interactions with breast cancer cells. **A)** Cell viability following doxorubicin treatment of SK-BR-3 WT and KO cells. The dose-response curve showed a comparable IC_50_ at 72 hours. **B)** MTT assay of SK-BR-3 WT and KO cells after 72 hours of incubation with doxo-EVs (60 nM of secreted doxorubicin). Doxo-EVs were isolated by differential ultracentrifugation from transfected and doxorubicin-treated HEK293T and subsequently analyzed by UHPLC-MS for doxorubicin quantification. Controls including doxorubicin alone are reported in the graph on the right. Graphs show Mean and SD of three biological replicates. **C)** Schematic representation of primers tested in ddPCR to detect the recombinant transcripts in EV-RNAs. The histogram below shows the observed transcript copy number per microliter in ddPCR experiments. Mean and SD refer to two independent experiments. **D)** cDNA synthetized from EV-RNA was amplified for detecting the recombinant transcripts in the full-length form (indicated by arrows) on agarose gel. **E)** Schematic representation of the *in vivo* study, from mice treatment (5 per condition) to recombinant EV-RNA detection by ddPCR in tumor xenografts. Mock doxo-EVs and antiHER2 doxo-EVs were tested. Subcutaneously injection was performed with a doxorubicin concentration of 0.5 µg/kg at tumor implantation site on day 0 and day 2. Image was created with BioRender.com. **F)** Representative agarose gel of the RNA quality (18S and 28S rRNAs, 2 μg RNA/well) obtained from tumor xenografts. The graph on the right shows the copies/µl of recombinant transcripts (each dot corresponds to one animal).

### Recombinant EVs reach breast orthotopic tumors and share their RNA cargo

Since doxorubicin experiments represent indirect evidence of cargo release, we prepared doxo-EVs from HEK293T cells for an *in vivo* study to detect horizontal delivery. The vesicular RNA cargo represents one of the best sources to be combined with sensitive technologies such as digital droplet PCR (ddPCR)(Pasini et al., 2021), (Notarangelo et al., 2019b). To understand whether recombinant EVs harbored the corresponding mRNAs, we designed a ddPCR assay to specifically detect relevant transcript fragments, which were indeed detected with different primer sets (**Figure 5C**). Interestingly, the whole chimeric sequences encoding the fusion proteins were amplified by PCR from the EV-RNA and showed the expected full-length on agarose gel (**Figure 5D**). We then moved to the *in vivo* study involving female athymic nude mice bearing JIMT-1 human breast tumor xenografts. JIMT-1 cells are trastuzumab-resistant(Zsebik et al., 2006) and represented a first choice to avoid HER2-mediated cytotoxicity and specifically detect horizontally transferred molecules. Mice were injected twice with doxo-EVs at the same doxorubicin concentration (0.5 µg/kg), on day one and day three, to be then sacrificed on day four (**Figure 5E**). As expected from the relative low dosage of doxorubicin administered, no statistically significant variations were observed in animal weight and tumor volumes among the different treatments (**Supplementary Figure 5A-B**). Therefore, we analyzed the RNA content of *ex-vivo* tumor cells to verify a detectable cargo of recombinant EVs. We isolated total RNA from tumor cell xenografts and used this template to detect EV-derived fusion protein-encoding transcripts by ddPCR. Interestingly, antiHER2 fusion transcripts were detected in tumor xenografts from at least 2 out of 5 animals, demonstrating a direct transfer from heterologous EVs *in vivo* (**Figure 5F**).

## Discussion

Several studies have reported tetraspanins as valuable tools for engineering and tracing cell-secreted EVs with fluorescent markers (Suetsugu, 2013), (Yoshimura et al., 2016), (Mathieu et al., 2021). Specifically, CD81 was fused with fluorescent reporters (Stickney et al., 2016), (Zuppone et al., 2023) or used as a direct targeting moiety against human placental laminin (Vogt et al., 2021). In this work, CD81 fusion proteins were exploited to track the cell engagement of secreted, heterologous EV sub-populations. As mainly indicated by imaging flow cytometry, the bulk of detected fusion proteins was not 100% vesicular, therefore fusion proteins could be subjected to parallel routes of packaging, clustering, and secretion (Meldolesi, 2022). Nevertheless, the ectopic CD81 expression stimulated EV secretion and slightly influenced the size of recovered particles. Cryo-EM experiments confirmed this trend but also presented multi-layered vesicles. Multi-lamellarity was circumstantially described but not yet elucidated if it derives from sample isolation procedures or represents discrete formations with potential biological roles (Broad et al., 2023), (Koo et al., 2021). Since this parameter changed across the tested conditions and mainly characterized the Mock condition (∼30% of bulk EVs), we reasoned that this parameter could be influenced by the relative single-EV protein content and/or turnover of endo/exocytosis processes. Further research involving dedicated strategies of single EV analysis, perhaps including tomography, is needed to address this aspect.

Seminal studies have already addressed the orientation of the extracellular domain of tetraspanins, including CD81, on the plasma membrane, showing that exposed sub-regions can have different degrees of conservation and serve homo or hetero interactions with tetraspanins (Seigneuret et al., 2001). Proteins like CD63, CD9, and CD81 have been recently proposed as a scaffold for inner and outer membranes to display fluorescent reporters, assuming a cell-membrane-equivalent protein orientation. There is circumstantial evidence that SCAMP3 and CD9 transmembrane proteins might configure a reversed topology on EVs compared to the plasma membrane (Cvjetkovic et al., 2016), (Hochheimer et al., 2019), and this outcome could be influenced by a repeated release turnover or budding *versus* fusion processes (O’Brien et al., 2022), potentially generating mixed orientations. By competitive interaction assays, we demonstrated that at least a fraction of secreted EVs presented an outward topology responsible for the specific binding to the HER2 receptor. In perspective, sorting specific EV sub-populations could allow investigation of the chimeric protein enrichment with a desired topology and in connection with elicited functions in recipient cells.

Despite the presence of GFP virtually contributing to a sub-optimal HER2 receptor binding, we had a chance to monitor EV-cell interactions *in vitro*. The mechanism underlying the preferential recruitment of heterologous EVs is not well understood. The proteo-lipid composition on cell and EV membranes may influence EV internalization and the propensity to uptake EVs (van Niel et al., 2022). In our experiments, two distinct HER2-positive and triple-negative breast cancer cell lines showed the same cumulative recruitment upon exposure to recombinant EVs for a few hours. Therefore, different interactions could result from intrinsic cell properties affecting the EV distribution *in vitro*. However, this observation needs to be evaluated in parallel with the positive engagement of recombinant EVs in HER2+ isogenic cells dictated by Trastuzumab light chains in our fusion protein. These experiments indirectly demonstrate that EV-cell interactions are contributed by different mechanisms possibly involving either proteins, lipids, or eventually nucleic acids. Notably, antiHER2 EVs were generally more prone to interact than CD81-GFP EVs, an occurrence possibly associated with a slight increase in diameter and multi-lamellar fraction (Figure 2). The existence of multiple mechanisms of EV-cell interactions can significantly limit the expectations from engineering approaches based on single peptides to increase targeting capabilities. A comprehensive characterization of the surfaceome (Meng et al., 2023), (Buzás et al., 2018), possibly including glycans (Goncalves et al., 2022), of a desired cell population could help customize a multi-modal targeting with enhanced specificity.

Once interacting during the time frame analyzed, live imaging acquisitions indicated a partial co-localization of green spots with lysosomes, suggesting that EVs may have different trafficking rates or follow non-degradative routes within recipient cells (Joshi et al., 2020), (Herrmann et al., 2021). Albeit the characterization of drug encapsulation efficiency and release kinetics need to be implemented and controlled, our results confirm the superior cytotoxic effects of secreted doxorubicin-EVs on HER2-positive breast cancer cells compared to the free drug, as already observed in other cell types (Farhat et al., 2022). This aspect would represent an advantage *in vivo* for reducing the effective drug dose and limiting off-targets. In line with cell-interaction assays, the cell-killing activity of recombinant doxorubicin-EVs was significantly mitigated upon removing the cell surface receptor, suggesting that a significant fraction of EVs can have a predictable outcome upon internalization. Therefore, the step of EV-cell interaction represents a priority to be addressed for targeted delivery. We also provide a direct indication through the RNA cargo horizontally shared *in vivo* and sensitively detected by ddPCR assays. This approach could be potentially applied to quantify recombinant RNA in tissue sections and address the penetration capacity of EVs.

In summary, this work prioritizes the profiling of both vesicular and cellular surfaceomes for a better comprehension of EV-cell interactions and a more efficient design and purification of EV-based nanovehicles.

## Acknowledgements

We warmly thank Dr. Deniz Ugurlar and Dr. Boxue Ma (ThermoFisher), who performed Cryo-EM image acquisitions.

## Author contributions

VGD conceived the study and supervised experiments; EG performed experiments and statistical analyses; GC contributed to ddPCR assays; MR and MP contributed to imaging and analysis; DP and RB contributed to doxorubicin detection; CT and MN participated in plasmid cloning and EV isolation; WQH and TTH supervised EV-cell imaging interaction; AC participated in starting EV isolation experiments. EG, TTH, and VGD wrote and revised the manuscript. All the authors contributed to manuscript writing and revision.

## Funding sources

The project has been supported by Fondazione Cassa di Risparmio Trento e Rovereto, Caritro (to VGD), and intramural resources (to VGD). The HTS, AICF, and MS Core Facility of CIBIO Department are supported by the European Regional Development Fund (ERDF) 2014-2020.

## Declarations

## Conflict of interest

The authors declare no conflict of interests.

## Ethics approval and consent to participate

Not applicable.

## Consent for publication

All authors agree for publication.

## Availability of data and material

Data supporting the findings are included in this article and the Supplemental information. The raw data and experimental materials are available from the corresponding author upon reasonable request.

## Notes

### Competing Interest Statement

The authors have declared no competing interest.

