## Supplementary figures for "CD81-guided heterologous EVs present heterogeneous interactions with breast cancer cells"

**Supplementary Figures and Figure legends**

**Supplementary Figure 1**

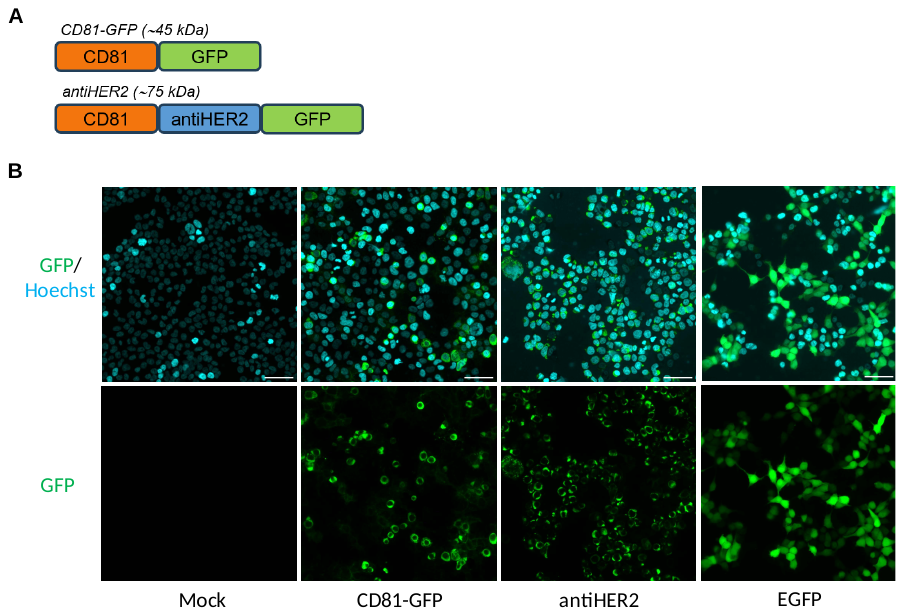

**A)** CD81-GFP and CD81-antiHER2-GFP (“antiHER2”) fusion proteins are depicted, with light chains of trastuzumab (antiHER2 moiety, blue) and turboGFP (tGFP, green). **B)** Spinning Disc confocal microscope to visualize the expression and localization of green-fluorescent fusion proteins and nuclei (Hoechst) in HEK293T cells after 48 hr of transfection. Mock and EGFP conditions are reported as negative and positive controls, respectively, of cell body staining. Scale bars correspond to 50 μm.

**Supplementary Figure 2**

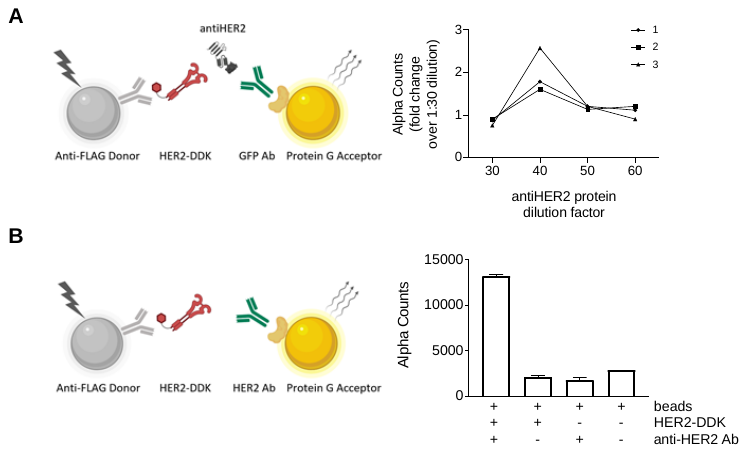

**A)** Sandwich designed for AlphaLISA assay to detect the binding of an *in vitro* translated antiHER2 protein to a recombinant HER2 (DDK-/FLAG-tagged). Alpha Count fold change shows binding specificity at dilution 1:40 of antiHER2 protein in three independent experiments (1-3). **B)** AlphaLISA competitive assay validation (see also Figure 3C). As shown in the graph, high Alpha Counts were obtained only in the presence of all the sandwich components.

**Supplementary Figure 3**

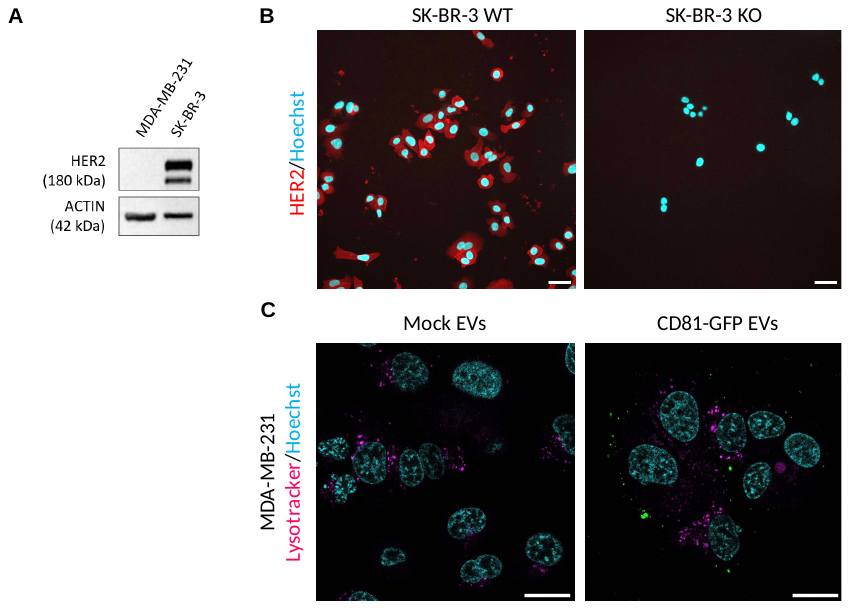

**A)** Immunoblotting to check HER2 expression in MDA-MB-231 and SK-BR-3 cells. **B)** Representative confocal images to show abrogation of HER2 expression in SK-BR-3 KO cells. The scale bar indicates 50 μm. **C)** Mock EVs were tested as negative control compared to GFP-positive EVs for EV uptake acquisitions as in Figure 4C. A ratio of about 30,000 bulk EVs per seeded cell was maintained to ensure GFP detection. GFP-EVs are shown in green, lysosomes are shown in magenta (Lysotracker red), and nuclei in cyan (Hoechst). The indicated scale bar corresponds to 10 μm.

**Supplementary Figure 4**

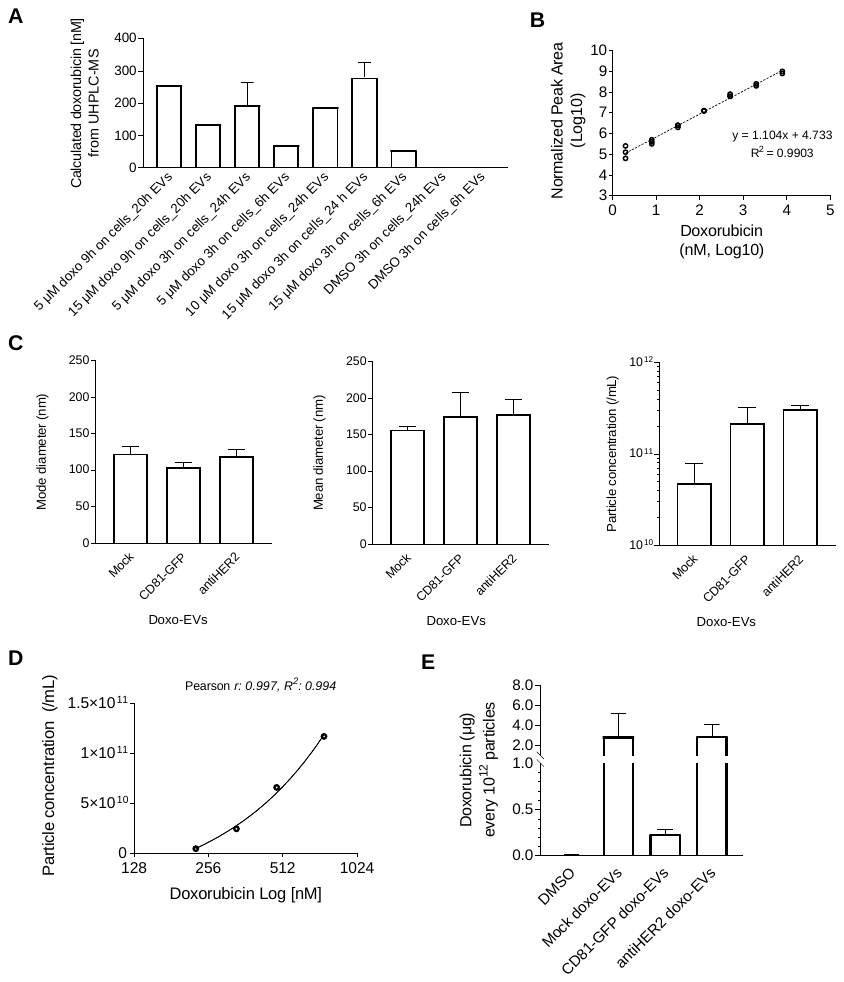

**A)** Dose (5, 10 or 15 μM), incubation time (3 or 9 hr), and release time (6, 20 or 24 hr) of doxorubicin secreted by transfected HEK293T cells. The Y axis indicates the drug concentration retrieved from isolated EVs by UHPLC-MS (see Methods). **B)** Calibration curve of doxorubicin standards run in parallel to doxo-EVs samples (see Methods). **C)** NTA analysis of doxo-EVs (mode and mean diameters, and particle concentration). **D)** Correlation between doxorubicin concentration retrieved from doxo-EV samples at UHPLC-MS and particle concentration from NTA measurements. **E)** Calculated doxorubicin relative abundance in doxo-EVs, normalized to particle number retrieved from NTA analysis. Mean and SD refer to two independent experiments.

**Supplementary Figure 5**

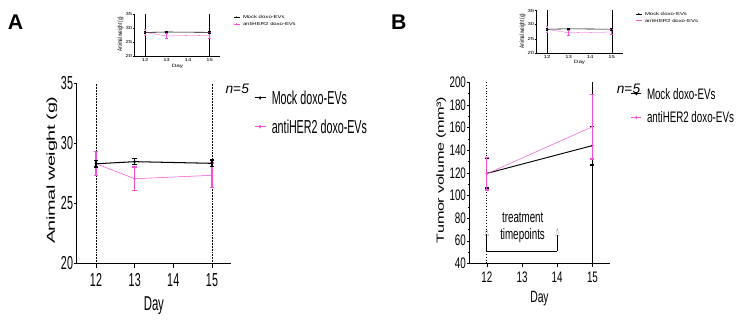

**A)** Animal weight measured during the *in vivo* study at day 12, 13 and 15. Mean and SEM are reported in the graph. **B)** Tumor volume measured before and after mice treatment (day 12 and 15). Mean and SEM are reported in the graph.
